# Assessment of anesthesia on physiological stability and bold signal reliability during visual or acoustic stimulation in the cat

**DOI:** 10.1101/827949

**Authors:** Alexandra Levine, Benson Li, Paisley Barnes, Stephen G. Lomber, Blake E. Butler

## Abstract

**Background:** Neuroimaging methods including fMRI provide powerful tools to observe whole-brain functional networks. This is particularly powerful in animal models, allowing these networks to be probed using complementary methods. However, most animals must be anesthetized for neuroimaging, giving rise to complications resulting from anesthetic effects on the animal’s physiological and neurological functions. For example, an established protocol for feline neuroimaging involves co-administration of ketamine and isoflurane – the latter of which is known to suppress cortical function.

**New Method:** Here, we compare this established protocol to alfaxalone, a single-agent anesthetic for functional neuroimaging. We first compare the two in a controlled environment to assess relative safety and to measure physiological stability over an extended time window. We then compare patterns of auditory and visually-evoked activity measured at 7T to assess mean signal strength and between-subjects signal variability.

**Results in Comparison with Existing Methods:** We show that alfaxalone results in more stable respiratory rates over the 120 minutes testing period, with evidence of smaller between measurements variability within this time window, when compared to ketamine plus isoflurane. Moreover, we demonstrate that both agents evoke similar mean BOLD signals across animals, but that alfaxalone elicits more consistent BOLD activity in response to sound stimuli across all ROIs observed.

**Conclusions:** Alfaxalone is observed to be more physiologically stable, evoking a more consistent BOLD signal across animals than the co-administration of ketamine and isoflurane. Thus, an alfaxalone-based protocol may represent a better approach for neuroimaging in animal models requiring anesthesia.

## 1. Introduction

Experiments in animal models continue to be critical for understanding the structure and function of the brain. For decades, cats have been successfully used as an animal model to study sensory systems, largely due to the remarkable similarities that they share with human cell types, neural pathways, and cytoarchitecture (Blake, 1979). Electrophysiological (Hubel and Wiesel, 1962; Liu et al., 2010), neuroanatomical (Lomber et al., 1995; Wong et al., 2015; Butler et al., 2016), behavioral (Heffner and Heffner, 1988; Wong et al., 2018), and imaging studies (Brown et al., 2014; Butler et al., 2015; Stolzberg et al., 2018) undertaken in the cat have played a critical role in advancing our knowledge of neural processing within visual and auditory systems, and the interactions between the two.

Recently, there has been increased interest in non-invasive neuroimaging methods like functional magnetic resonance imaging (fMRI) for the study of sensory system function in animal models. This approach offers several advantages, including the ability to examine perception at the whole-brain level, and the ability to undertake longitudinal, within-animal studies of sensory system development (which in turn aids in reducing the number of animals required to power meaningful comparisons). This offers a distinct advantage over other methods such as electrophysiological studies, which are highly invasive and are limited in their capacity to evaluate processes occurring over spatially disparate neural networks. fMRI measures changes in the ratio of oxygenated to deoxygenated blood, or blood-oxygen-level-dependent (BOLD) signals (Ogawa et al., 1990). An increase in this BOLD signal is thought to reflect increased neuronal activity compared to a baseline measurement (Buxton and Frank, 1997; Logothetis et al., 2001; Ferris et al., 2006). Moreover, by measuring the temporal coherence of the BOLD signal across spatially disparate areas of the brain, it is possible to estimate the degree to which these areas are functionally connected into networks that support perception and associated behaviors.

Modern fMRI scanners can provide spatial resolution with 1 mm precision, but the accuracy of these measures depends critically on minimizing subject movement within the scanner. While most human participants can be instructed to remain still during imaging sessions, this is not possible in most other animals, and meaningful measurements must thus be taken under anesthesia. Anesthetic agents help minimize potential stress and fluctuations in behavior that can affect the quality of data retrieved. However, the use of anesthetics during fMRI necessitates due consideration be given to the effects of the drugs themselves (Ueki et al., 1992; Biermann et al., 2012), as these agents can influence neurovasculature by changing cerebral blood flow, blood volume, and rate of oxygen metabolism (Gao et al., 2017), and can suppress neuronal activity by reducing excitatory synaptic transmission or increasing inhibitory transmission (Richards, 1983). Further complications may include physiological variability based on the drug type, concentration, and route of administration (Peng et al., 2010; Nagore et al., 2013; Aksenov et al., 2015; Ros et al., 2017). Therefore, there is a need to establish a robust and reliable anesthetic protocol that will facilitate bridging the gap between animal and human neuronal organization and function.

Fortunately, decades of electrophysiological work in the cat has revealed a great deal about the effects of different anesthetic agents on recorded neural function. A common protocol involves the continuous infusion of ketamine alongside other agents such as sodium pentobarbital, xylazine, or diazepam to induce and maintain anesthesia (e.g. Heil and Irvine, 1998; Miller et al., 2002; Pienkowski and Eggermont, 2009). This protocol has evolved over time; for example, early studies found that ketamine infusion reduces spontaneous and peak firing rates in auditory cortex (Zurita et al., 1994), possibly due to altered sensory perception and reduced cortical glucose metabolism (Crosby et al., 1982; Oye et al., 1992). To reduce the amount of ketamine required for anesthesia (and reduce these suppressive effects, accordingly), Jezzard et al. (1997) proposed to pre-medicate with ketamine but maintain sedation with isoflurane (1-2%, gas). However, other studies showed that isoflurane redirected cerebral blood flow, and reduced neural activity recorded in various visual brain regions by up to 50% (Harel et al., 2002; Olman et al., 2003; White & Alkire, 2002). Isoflurane has recently been shown to suppress resting-state connectivity in the primary somatosensory cortex of non-human primates as well (Wu et al., 2016). Therefore, when establishing the protocol for initial fMRI experiments in the cat, Brown et al. (2013) reduced isoflurane concentrations to the minimum level required to maintain sedation (0.4-0.5%) and supplemented with a continuous rate infusion of ketamine (0.6-0.75 mg/kg/hr). Under this protocol, the authors were able to record BOLD signal changes up to 6% in some but not all auditory regions. The protocol was used in subsequent auditory-evoked studies (Hall et al., 2014; Butler et al., 2015) and in an examination of resting-state connectivity using fMRI (Stolzberg et al., 2018).

In spite of successive revisions, the combination of isoflurane and ketamine is known to result in widespread cortical deactivation in other animals (e.g. Hodkinson et al., 2012). Moreover, these effects appear to differ by brain region/sensory modality such that the co-administration of ketamine and isoflurane may limit the ability to study sensory processes beyond audition (Oye et al., 1992; Ries & Puil, 1999; Hoflich et al., 2017). Thus, there remains a need to develop an anesthetic protocol that can induce and maintain a light anesthetic plane sufficient to suppress movement, without drastic reductions in cortical activity across multiple brain regions.

Several potential agents were considered in the current study with important limitations in mind. In addition to the complications related to isoflurane described above, ketamine is known to be a dissociative agent, disrupting the central nervous system and causing a cataleptic state with dose-dependent hallucinations; thus, some protocols in routine use (e.g. ketamine plus diazepam/midazolam/xylazine) were deemed suboptimal for sensory-evoked neuroimaging. Moreover, a single-agent approach was considered practical in order to avoid complications inherent to maintaining a stable level of anesthesia during dynamic and complex drug interactions (an assumption critical to interpreting neuroimaging data averaged over extended time periods). In addition, many agents were excluded because their mechanism of action was deemed not conducive for measuring BOLD signals (Table 1). Others were eliminated in consultation with veterinary care staff due to concerns with respect to physiological effects. As a result, alfaxalone was considered to be the strongest candidate protocol to serve as an alternative to the coadministration of ketamine and isoflurane as a primary anesthetic agent for fMRI. Alfaxalone: i) has dose-dependent effects on cardiovasculature, respiration, neuronal activity, and neuromusculature (Warne et al., 2015; Whittem et al., 2008; Muir et al., 2009; Taboada and Murison, 2010; Baldy-Moulinier et al., 1975) which may allow for more predictable changes in BOLD response; ii) has been found to sufficiently maintain a stable level of anesthesia for up to 2-hours as a stand-alone agent (Tamura et al., 2015; Deutsch et al., 2017) which in our experience is the typical amount of time required for neuroimaging in cats; and iii) has been successfully administered intravenously to maintain anesthesia in cats during surgical procedures in our own laboratory, and by others (Beths et al., 2014; Nagakubo et al., 2017) with minimal physiological side effects.

**Table 1.**
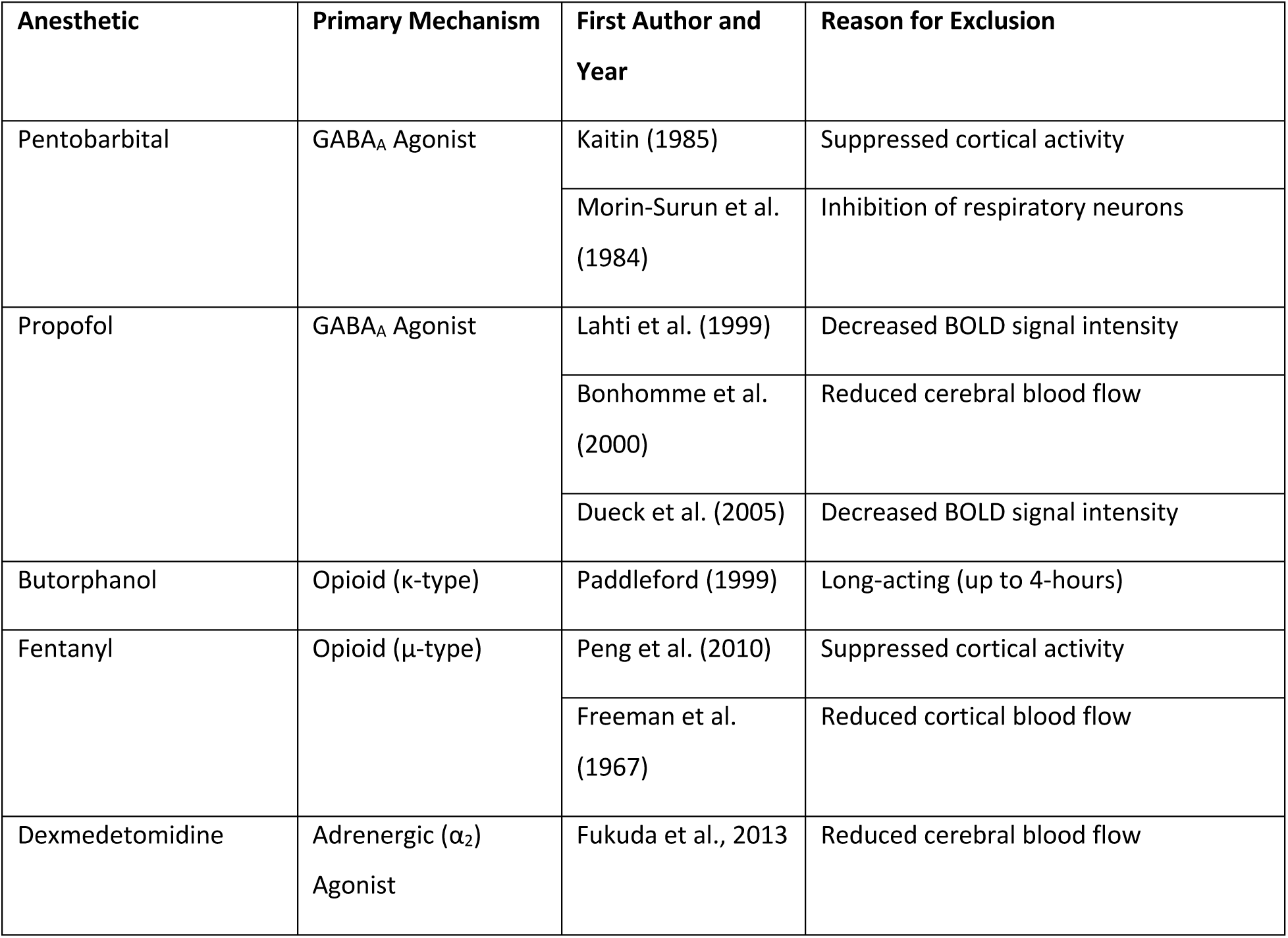
Studies examining cardiovascular, respiratory, and neural effects of anesthetic agents.

Here, we compare an alfaxalone (Alf) protocol with a previously established protocol consisting of a combination of isoflurane and ketamine (Iso+Ket), providing detailed measures of physiological stability as well as evoked activity in cats during fMRI. The investigation is separated into two parts; in the first study, we evaluate the physiological stability of each protocol in an operating suite. A light anesthetic plane was maintained for a minimum of 2-hours while cats were exposed to mock scanner noises at 90 dB and vital signs (heart rate, respiratory rate, end-tidal CO_2_, blood pressure, etc.) were recorded. In the second study, both protocols were employed in the scanner while BOLD signal changes were recorded in response to visual and auditory stimuli. This study is the first to evaluate different anesthetic protocols in cats by directly comparing BOLD signal responses. Quantification of these signals will provide insight into the contribution of different anesthetics on neural activity, and potentially offer an alternative option to the combination of isoflurane and ketamine.

## 2. Methods

### 2.1 Animals

Two healthy adult domestic short-hair cats were used in study one, and a total of 12 cats were compared in the second study. Animals were born to pregnant queens obtained from a commercial laboratory animal breeding facility (Liberty Labs, Waverly, NY), and were housed as a clowder. Normal hearing status was confirmed at approximately 3 months of age using auditory brainstem responses. All procedures were conducted in accordance with the Canadian Council on Animal Care’s Guide to the Care and Use of Experimental Animals and were approved by the University of Western Ontario Animal Use Subcommittee of the University Council on Animal Care.

### 2.2 Study 1: Physiological Stability during Anesthesia

#### 2.2.1 Anesthesia

The first study was conducted in a surgical suite in order to evaluate the safety and stability of the selected protocols. In the Alf protocol, the animal was first pre-medicated with dexdomitor (0.04 mg/kg, i.m.) prior to catheter placement. Sedation was confirmed after 10 minutes by the absence of a paw-pinch reflex. Ophthalmic ointment was applied to prevent drying of the eyes, body temperature was maintained at 37°C using a circulating warm water pad, and an indwelling 22g catheter was placed in the cephalic vein to facilitate maintenance of anesthesia. A bolus dose of alfaxalone (0.3-0.5 ml, i.v.) was administered to achieve deeper anesthesia and the animal’s larynx was sprayed with xylocaine prior to intubation. The animal was placed in a sternal position on the surgical table, and anesthesia was maintained through continuous infusion of alfaxalone (7 mg/kg/hr, i.v.), while 100% oxygen was provided at a rate of 1.0L/min. Finally, a bolus dose of atipamezole (0.27 ml, i.m.) was administered to reverse any residual effects of the dexdomitor.

For the Iso+Ket protocol, animals were pre-medicated with a combination of dexdomitor (0.022 mg/kg, i.m.), ketamine (4 mg/kg, i.m.), and acepromazine (0.05 mg/kg, i.m.). Sedation was confirmed, the animal’s core temperature was maintained, ophthalmic ointment was applied, and a catheter was placed for anesthetic maintenance as above. The animal was placed in a sternal position on the surgical table, and a continuous infusion of ketamine (5 ml/kg/hr, i.v.), combined with gaseous isoflurane (0.5% in oxygen provided at a rate of 1.0L/min) was used to maintain anesthesia. The reversal of dexdomitor was not necessary in this protocol as premed volume was lower and consequently would not be expected to have effects lasting into the experimental period. Approximately 60 minutes into the session, the rate of ketamine infusion was increased to 6.25 ml/kg/hr (i.v.) and isoflurane was reduced to 0.25% in order to mimic the protocol developed previously for imaging, in which these changes are required prior to functional image acquisition to optimize BOLD signal.

At the end of each session, anesthesia was discontinued, and animals were monitored until they recovered fully from anesthetic effects. Animals anesthetized with alfaxalone received a bolus dose of butorphanol (0.2 mg/kg, s.c.; opioid analgesic) to counteract hyperkinesia, a side-effect commonly observed during post-anesthetic recovery from prolonged IV administration of alfaxalone in cats (Whittem et al., 2008). The intubation tube was removed when the animal exhibited a gag reflex and increased jaw tone, and following recovery, the indwelling catheter was removed and the animal was returned to their clowder. Each agent was tested twice in each animal for a total of 4 sessions per agent.

#### 2.2.2 Data Recording

To mimic conditions in the scanner, the animal was presented with previously recorded scanner noise through foam insert earbuds (Sensimetric S14) at 90 dB SPL for the duration of experimental sessions. Each agent’s ability to induce anesthesia, maintain a lightly sedated state for 2-hours, and to allow for uneventful recovery was noted. Anesthetic and physiological stability was evaluated by monitoring and recording parameters including autonomic reflexes (e.g. paw-pinch, gag, palpebral) and vital signs (e.g. heart rate, end-tidal CO_2_, respiratory rate, peripheral capillary oxygen saturation, blood pressure, and mean arterial pressure) in 5-minute intervals.

### 2.3 Study 2: fMRI

The second study sought to compare the auditory- and visually-evoked BOLD signals recorded while animals were anesthetized with each candidate agent. A group of 6 cats were scanned while anesthetized with alfaxalone, and results were compared to a group of 6 sex- and age-matched animals scanned previously using the exact same equipment and experimental procedure.

#### 2.3.1 Animal Preparation and Anesthesia

For both the Iso+Ket and Alf protocols, anesthesia was induced and maintained as described for Study 1 above. Once anesthetized, the animal was placed in a sternal position within a custom-built Plexiglass sled. Phenylephrine hydrochloride and atropine sulfate ophthalmic solutions were applied to both eyes to dilate the pupils and retract the nictitating membranes. Lubricated contact lenses were placed in both eyes (a blackout lens in the left eye, and a clear lens in the right eye). This permitted visual stimuli to be brought into focus on the retina and have visual signals preferentially sent to the left hemisphere. MRI-compatible foam insert earphones (Sensitmetrics S14) were inserted in each ear to allow for the presentation of auditory stimuli, and the animal’s head was stabilized within a custom 8-channel radio-frequency (RF) coil. Vital signs (heart rate, respiratory rate, end-tidal CO_2_, inspiratory CO_2_, percent oxygen saturation, systolic/diastolic and mean blood pressure, and rectal body temperature) were monitored throughout the scanning session. At the conclusion of the imaging session, anesthesia was discontinued and animals were recovered as outlined in Study 1 above.

#### 2.3.2 Stimuli

Visual stimuli were generated with PsychoPy (Peirce, 2007; 2009) and presented through a Dell laptop to an Avotec SV-6011 Rear Projector. From their sternal position within the bore of the magnet, the animals eyes were located approximately 75 cm from an acrylic screen (H = 14.5 cm, W = 19cm), which was viewed through a custom-built mirrored periscope. The stimulus extended 14.5 visual degrees horizontally and 11 degrees vertically and consisted of a black and white flickering checkerboard (8 ring-segments of 16 wedges) on a grey background (100% luminance contrast, 50% luminance background), counter-phase flickering at 5Hz. The stimulus was arranged in a simple ON/OFF block design, where the OFF block consisted of a blank grey screen, each block lasting 30s. The animal’s gaze was assessed visually through the scanner bore before the acrylic screen was placed at the end of the bore.

Auditory stimuli were generated using Audacity® recording and editing software (Audacity Team, 2019), and consisted of a 30s stimulus consisting of 400ms broadband noises separated by 100ms silent gaps. This stimulus was arranged in an ON/OFF block design, where the OFF block consisted of a 30s period of silence. Sounds were presented diotically from a Dell laptop through an external Roland Corporation soundcard (24-bit/96 kHz; Model UA-25EX), a PylePro power amplifier (Model PCAU11), and Sensimetrics MRI-compatible ear inserts (Model S14). All stimuli were calibrated to 80-90 dB SPL using an ear simulator (Bruel & Kjaer, Model # 4157).

#### 2.3.3 Scanning Parameters

Data were collected using an ultra-high-field 7T Siemens MRI human head-only scanner located at the Centre for Functional and Metabolic Mapping at the Robarts Research Institute operating at a 350 mT/m/s slew rate. An automated 3D mapping procedure (Klassen & Menon, 2004) was used to optimize the magnetic field (B_0_ shimming) over the specific volume of interest.

High-resolution structural T1-weighted MP2RAGE images were acquired prior to functional scanning with the following parameters: isotropic voxel size of 0.5mm^3^, 80 slices, FoV=96mm, TE=3.95ms, TR= 6000ms, TI =800ms, and a flip angle of 4. Functional images were acquired over the whole brain in axial orientation with a single shot echo-planar imaging (EPI) acquisition with grappa acceleration and the following parameters: isotropic voxels 1mm^3^, 38 slices (interleaved), FoV=72mm, TE=22.0ms, TR= 2000ms and a flip angle of 60 degrees. Each functional scan (visual- and auditory-evoked) lasted six minutes, and consisted of alternating 30 second blocks of stimulus and baseline conditions.

#### 2.3.4 Image Analysis

T1-weighted structural images were processed with a combined approach of automated and manual processing. The structural images were skull-stripped with use of MRIcron (NITRC; Rorden & Brett, 2000) and FSLmath functions (FMRIB’s toolbox, Oxford, UK; Smith et al., 2004, http://fsl.fmrib.ox.ac.uk/fsl/fslwiki/).

First-level statistical analysis of each animal’s functional data was carried out using FEAT processing in FSL (Woolrich et al., 2001). The functional images were skull stripped using FSL’s Brain Extraction Tool (BET). Preprocessing began with the removal of the first 2 volumes (4s) of the scan to allow for the scanner magnetic field to stabilize and reach magnetic saturation.

The following were also applied: motion correction (MCFLIRT; though the movement is nearly non-existent during these procedures), spatial smoothing (Gaussian, FWHM, 2mm) and a temporal high-pass filter cut off (0.01Hz/60s). First-level general linear model analysis (FILM) was then carried out, where regressors for each condition-block were convolved with a gamma hemodynamic response function (phase = 0, standard deviation = 3s, mean lag = 6s; the BOLD signal time course in cats has been shown to closely resemble that observed in humans and non-human primates [Brown et al., 2013]).

Each individual EPI sequence underwent time series pre-whitening (Smith et al., 2004), allowing us to carry through contrasts for higher-level analysis to test for group effects; individual animal GLM results were co-registered to the coordinate space of the high-resolution structural image for each participant using FMRIBs Linear Image Registration Tool (FLIRT; Jenkinson et al., 2002). Further analysis compared differences in average BOLD signal change under each anesthetic agent across all voxels within a given region of interest. Visual ROIs included primary visual cortical areas 17, 18 and 19, as well as the lateral geniculate nuclei (LGN) of the thalamus. Auditory ROIs included primary (A1) and second (A2) auditory cortex and the medial geniculate nuclei (MGN) of the thalamus.

Mean BOLD signal changes (relative to baseline) evoked by visual and auditory stimuli were extracted for each ROI of each animal, using FEATquery (FMRIB toolbox in FSL). To carry this out, FEATquery takes each participant’s high-resolution structural scan and co-registers it to the feline template space (CATLAS, Stolzberg et al., 2017) using FLIRT multi-registration, in which all the predefined functional ROIs are defined (Fig. 1).

**Figure 1.**
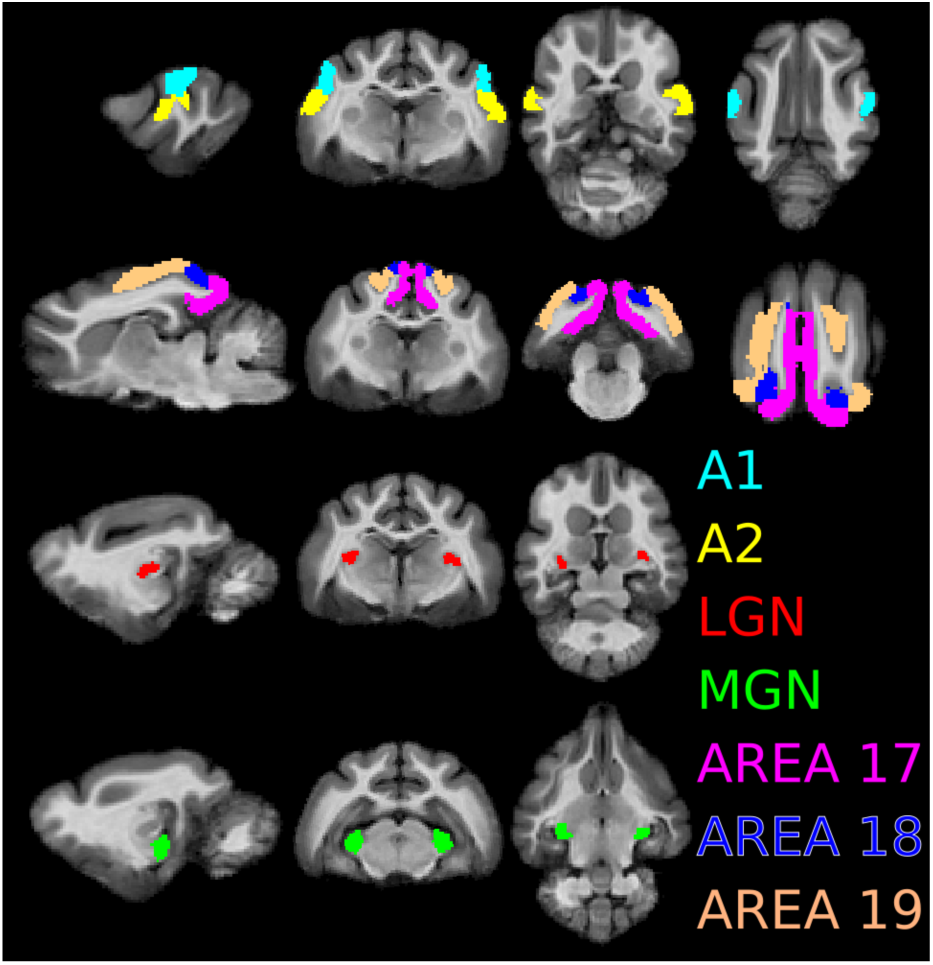
Visual and auditory regions of interest (ROIs) used in analysis displayed in feline template space (CATLAS; Stolzberg et al., 2017). Visual ROIs (LGN, 17/18/19) contralateral to the open eye were isolated for analysis.

## 3. Results

### 3.1 Study 1 - Physiology

Two animals were observed two times under each anesthetic protocol in the operating suite to evaluate the stability of heart and respiratory rates across anesthetics. These physiological measures are commonly yolked; in practice, respiratory rate is used as an indicator of anesthetic depth where high rates are more representative of an awake state (Myles, 2007). The data from the present evaluation, averaged across animals and runs for each anesthetic agent tested, are presented in Figure 2A. To examine the variability in heart and respiratory within a test session, the change in each measure between successive timepoints (i.e. the change in heart/respiratory rate across each 5-minute interval; values presented in Figure 2B) was calculated, and a Wilcoxon rank-sum test with continuity correction was conducted. On this time scale, changes in heart rate were not different across anesthetics (W = 3466.5, p = 0.0843); however, the respiratory rate differed significantly (W = 1937.5, p < 0.001) suggesting greater within-session stability under alfaxalone compared to the coadministration of ketamine and isoflurane.

**Figure 2.**
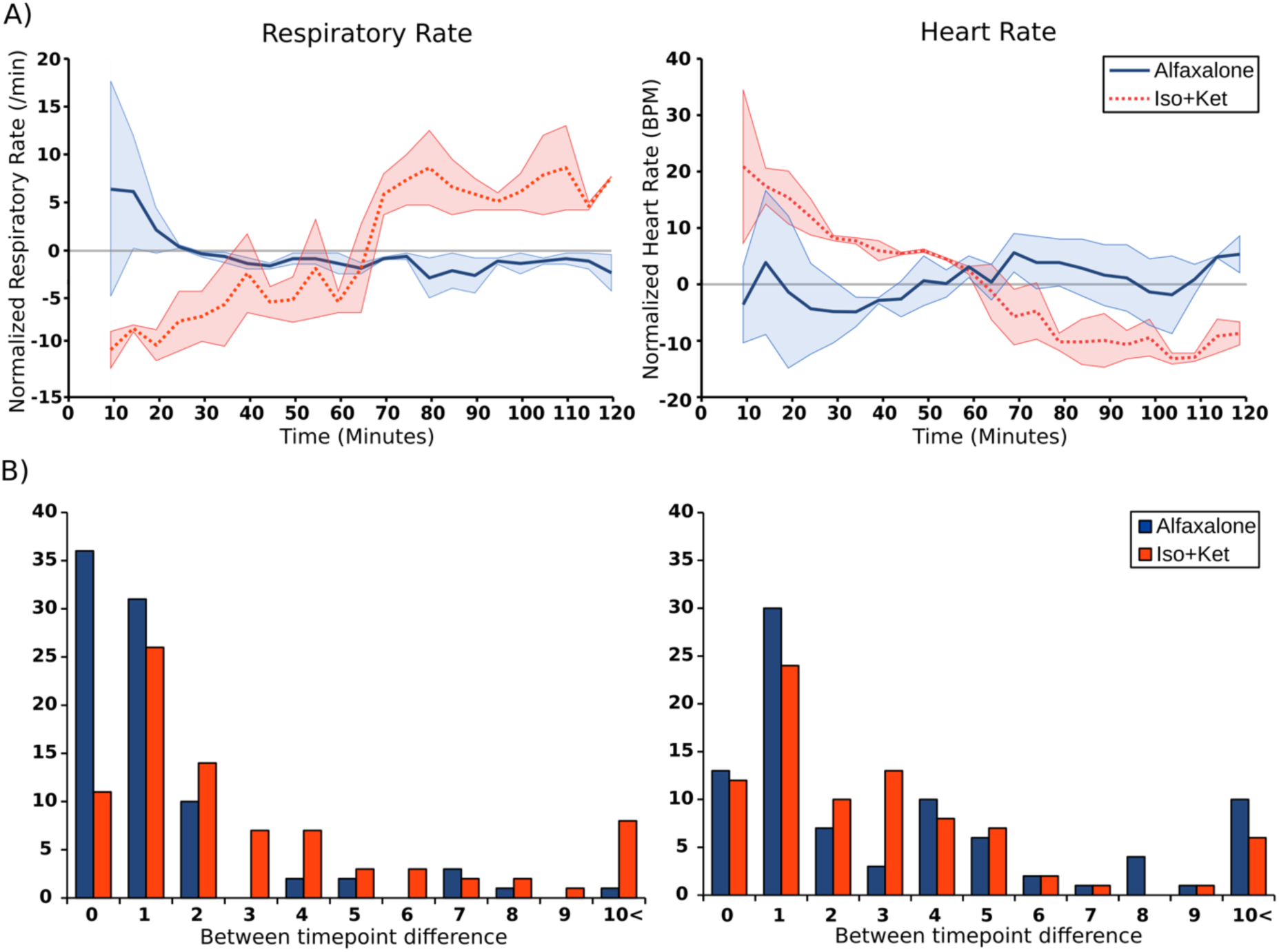
A) Normalized respiratory rates (breaths per min) and heart rates (beats per min) under alfaxalone and co-administered ketamine and isoflurane. Data were normalized to the mean rate for an individual run, and then normalized values were averaged across animals and runs. Shaded regions represent the standard error of the mean. The first 10 minutes of each measurement period were omitted from analysis to account for setup of monitoring/recording devices. B) A histogram representing change magnitude in successive measurements of respiratory and heart rate under each anesthetic tested. Lower values indicate relative stability in the variable measured, while larger values indicate increased variability over time.

It is important to note that increased variability under ketamine plus isolflurane is not entirely due to the drugs per se; imaging under this protocol requires that the rate of ketamine infusion be increased and the concentration of isoflurane reduced approximately 60 minutes into a testing session in order to acquire functional images with measurable BOLD signal (Brown et al., 2013; Stolzberg et al., 2018). As a result, physiological measures such as heart rate and respiratory rate often change dramatically at this point in time (observable in Figure 2A). As described above, these shifts indicate drift towards a lighter anesthetic plane, as is necessary to provide increased BOLD signal in the presence of isoflurane. However, this also increases the risk of the animal becoming alert, which should be avoided. In contrast, the rate of alfaxalone infusion can remain unchanged for the duration of the session, resulting in more stable vital signs throughout.

### 3.2 Study 2 - fMRI

#### 3.2.1 BOLD signal strength

To examine whether a global difference in BOLD signal amplitude exists between anesthetics, normalized percent BOLD signal changes (hereafter referred to as BOLD signals) were extracted from all selected ROIs, in each animal.

As an initial test, a mixed ANOVA was performed on these BOLD signals, where anesthetic protocol (Iso+Ket/Alf) was treated as a between-subjects factor, and within-subject factors included two stimulus types (auditory/visual) and seven ROIs (auditory cortical areas A1 and A2, the MGN, visual cortical areas 17, 18, and 19, and the LGN). A significant interaction was observed between stimulus type and ROI (*F*[11, 110] = 13.505, *p* < 0.001) demonstrating that across anesthetic protocols, the BOLD signal observed in a given ROI depended on the nature of the stimulus presented. The comparison of between-subjects effects across all regions and conditions revealed no significant effect of anesthetic protocol (*F*[1, 10] = 1.108, *p* = 0.337). Overall, these results indicate that, as expected, auditory and visual stimuli evoke different patterns of activity across sensory brain regions. Moreover, the patterns of evoked activity are similar across the anesthetics tested. While this suggests that both alfaxalone and co-administered ketamine and isoflurane may be appropriate for imaging experiments, this analysis provides little insight into the consistency and stability of the BOLD signal measured under each. We thus conducted planned post-hoc tests investigating the variability of these signals.

#### 3.2.2 BOLD signal variability

Figure 3 shows the BOLD signals recorded in each ROI broken down by stimulus type (auditory, visual) and anesthetic protocol for each animal tested. Doing so allows BOLD signal variability to be observed across all ROIs (bilateral MGN, A1, & A2; LGN, 17, 18, & 19 contralateral to the opened eye) and individual subjects. Across regions of interest, between-subjects variability in the BOLD signal evoked by stimuli to which a region is typically responsive (i.e. the signal recored in A1/A2 in response to sound) was greater under ketamine plus isoflurane than under alfaxalone. Further, signals recorded in response to stimuli to which an ROI is *not* typically responsive (i.e. the signal recorded in areas 17/18/19 in response to sound), were also far more variable under Iso+Ket. This latter analysis provides a measure of signal variability in the absence of evoked activity (i.e. a measure of background activity under a given anesthetic agent).

**Figure 3.**
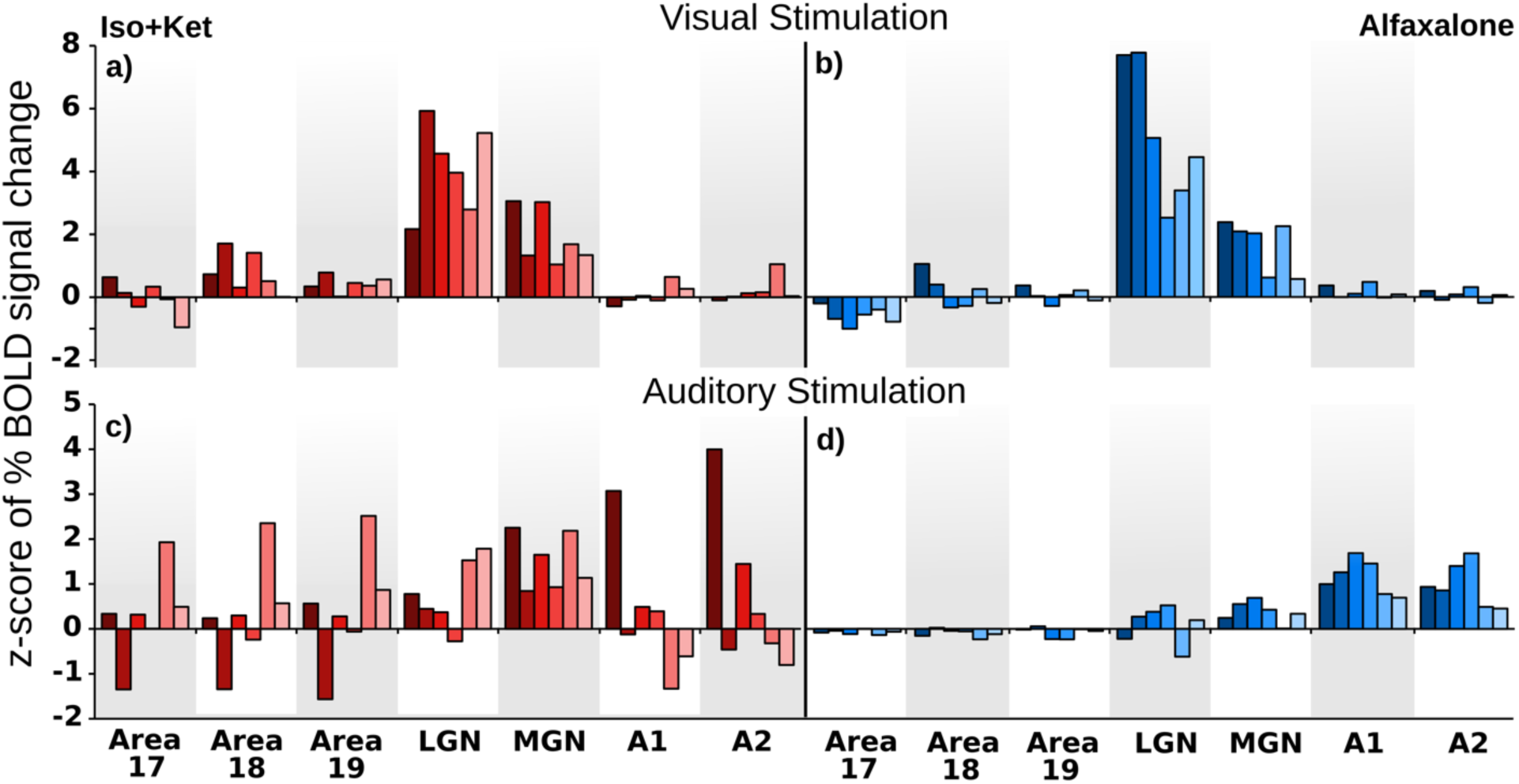
Individual z-scores of BOLD signal changes within all regions of interest in response to visual (a & b) and auditory (c & d) stimulation under coadministration of ketamine and isoflurane (a & c) or alfaxalone (b & d). Data are thresholded using clusters determined by Z>2.3 and a corrected cluster significance threshold of p=0.05 (Worsley, 2001).

To quantify variability in more detail, the range of the normalized BOLD signal across animals for each region of interest and anesthetic protocol is presented in Figure 4. With respect to visually-evoked activity, the between-subjects variability in BOLD activity was highly similar across all ROIs. In response to sound, the signal recorded was less variable under alfaxaolone than under co-administered ketamine and isolflurane across all ROIs. Thus, while both anesthetics evoke similar mean BOLD signal amplitudes (as evidenced by the absence of a statistically significant effect of anesthetic protocol, as described above), between-subjects variability is decreased under alfaxalone, notably in response to sounds.

**Figure 4.**
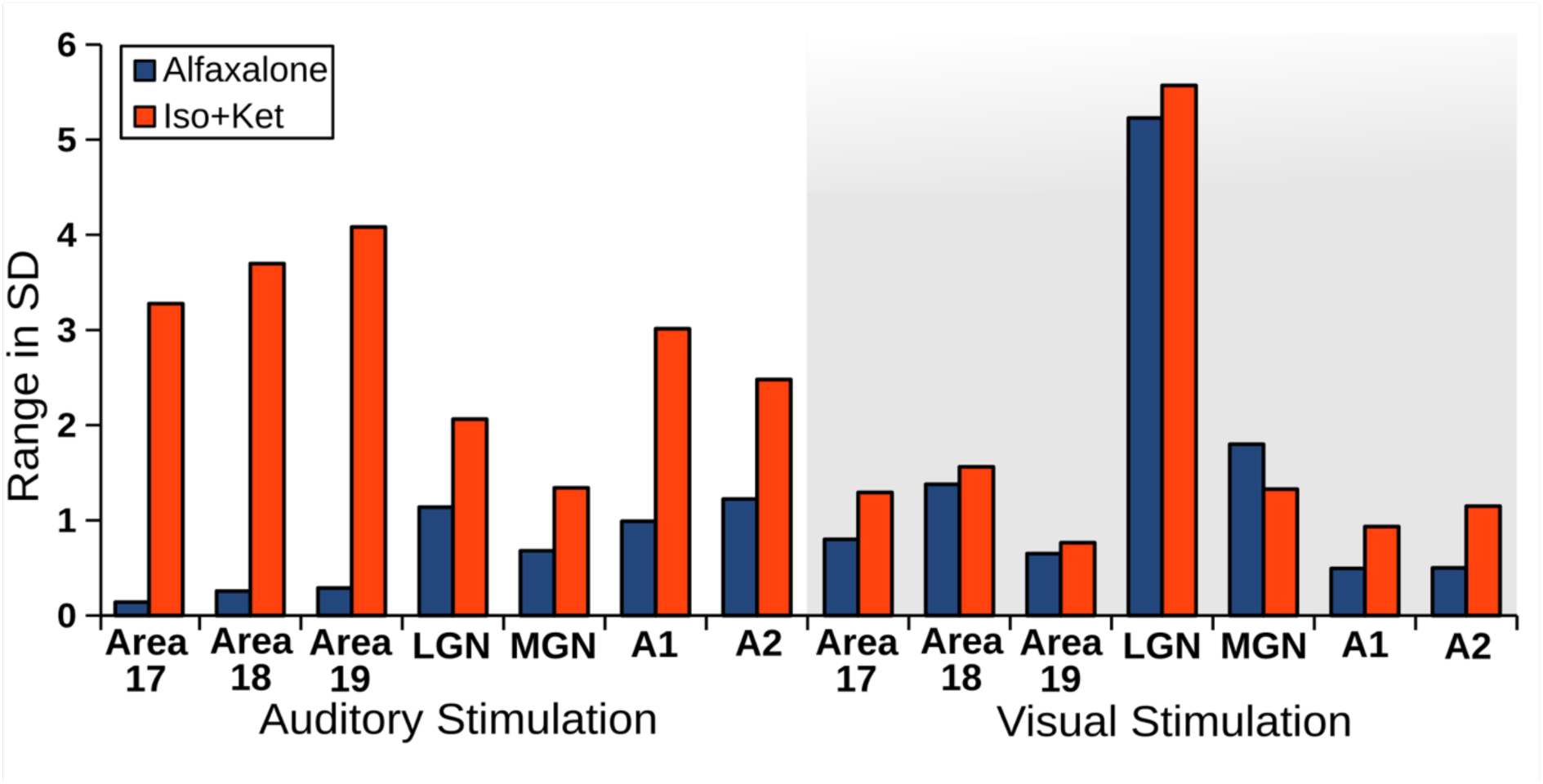
Range (max-min) of z-scored percent BOLD signal change across animals for each region of interest studied, in response to auditory and visual stimuli under alfaxalone (blue) and co-administered ketamine and isoflurane (red).

## 4. Discussion

This study was undertaken to compare and contrast potential anesthetic protocols for functional neuroimaging with a focus on 1) physiological stability and 2) optimizing BOLD signal across stimulus modalities. While all anesthetics affect neural function, their use remains necessary in most animal models to minimize movement and stress, and to ensure animal safety within the magnet. Thus, it is important to establish a regime that strikes a balance between achieving and maintaining safe and stable anesthesia, while optimizing cortical activity. Previous studies have used a variety of anesthetic agents in an attempt to optimize responses; one common protocol involves the co-administration of ketamine and isoflurane– an approach that has been shown to allow for the observation of sound-evoked BOLD activity (Brown et al., 2013; Hall et al., 2014; Butler et al., 2015; Stolzberg et al., 2018). However, this combination can produce physiological instability and has suppressive effects on cortical activity (Zurita et al., 1994; Harel et al., 2002; Olman et al., 2003; Hodkinson et al., 2012). It therefore remains important to explore alternative and perhaps more reliable protocols. Here, we provide physiological and neuroimaging evidence that supports the use of alfaxalone as a stable and consistent anesthetic agent for neuroimaging.

### 4.1 Anesthesia and Physiology

The primary goal of the physiological pilot described above, was to determine the safety and stability of candidate protocols using heart and respiratory rates as indicators. Importantly, these measures have been shown to reflect anesthetic depth (Musizza & Ribaric, 2010; Thomas & Lerche, 2011). Moreover, fluctuations in either heart or respiratory rate have been shown to interfere with neural signals measured by fMRI (Gao et al., 2017; Birn et al., 2003; Abbott et al., 2005; Kastrup et al., 1999). In part, the instability of co-administered ketamine and isoflurane reflects a simultaneous increase in ketamine infusion rate and decrease in isoflurane concentration approximately 60 minutes into an imaging session that is necessary to allow for BOLD signal visualization (Brown et al., 2013; Hall et al., 2014; Butler et al., 2015; Stolzberg et al., 2018). Thus, this previously established protocol includes a trade-off between measurable BOLD responses and increased risk of the animal becoming alert in the scanner. Conversely, the alfaxalone protocol described here maintains a constant infusion rate for the duration of the experimental session, and thus results in only small-scale changes in heart and respiratory rate over time, consistent with previous examinations of the anesthetic properties of alfaxalone for other applications (Beths et al., 2014; Muir et al., 2009). In addition to long-term stability, respiratory rate under alfaxalone was also shown to be more stable across shorter duration measurement intervals (5 min; Figure 2B). Both protocols examined attained safe levels of anesthetic depth for functional imaging; however, alfaxalone resulted in greater respiratory stability across individuals and sessions when compared to the co-administration of isoflurane and ketamine.

### 4.2 Anesthesia and BOLD

Having demonstrated the safety of both candidate protocols, the imaging experiment (Study 2) sought to compare and contrast BOLD signal changes evoked in auditory and visual thalamic and cortical regions of interest under each anesthetic. To the best of our knowledge, this is the first study to provide such a comparison. Here, we demonstrate that both protocols facilitate comparable mean levels of overall evoked BOLD activity across ROIs. These results are in accordance with similar mechanisms of action between alfaxalone and isoflurane (Lambert et al., 2003; Nakahiro et al., 1999). Examining these patterns of evoked activity in more detail, more reliable and consistent neural responses were observed under alfaxalone, evidenced by decreased BOLD signal variability across animals (Figures 3 & 4). Interestingly, this effect is most evident in sound-evoked activity, while patterns of BOLD activity evoked in response to visual stimuli are qualitatively similar across anesthetics in the current study. The reliability of recorded BOLD signals is highly important, as fMRI analyses often involve averaging or subtractive computations across blocks of data acquired over the duration of an imaging session. That the activity recorded under co-administered ketamine and isoflurane is highly variable within a given ROI means these between-block contrasts may be particularly susceptible to anesthetic effects. Interestingly, signals measured in brain regions not traditionally associated with a particular stimulus modality (e.g. BOLD signal estimates from primary visual cortex in response to auditory stimulation) were more variable under co-administered ketamine and isoflurane than under alfaxalone, suggesting signal variability associated with the former extends well beyond effects on stimulus-evoked activity. This activity may reflect between-subjects differences in non-selective suppression observed under isoflurane (Wu et al., 2016) or the potentially dissociative effects of ketamine (Abel et al., 2003) – either of which presents a challenge to the interpretation of stimulus-evoked signals. Substantial differences in evoked BOLD signal were observed in the current study across stimulus type. For example, greater thalamic activity was observed in response to visual stimuli than to sound. This could reflect a number of underlying causes, including differences in the extent to which the visual stimulus (a flashing, whole-field checkerboard) and auditory stimulus (white noise bursts) evoke robust activity within their respective ascending pathways. Importantly, while the visual stimulus evoked a greater BOLD signal within the presumptive auditory nucleus of the thalamus (MGN) than did sound, the results suggest that only the sound-evoked activity resulted in cortical activation. Thus, while visually-evoked activity in MGN could reflect crossmodal inputs, or spatial spread of the robust activity recorded in the nearby LGN, only sound-evoked thalamic signals appear to be faithfully translated to auditory cortical activity. Increased consistency between individual animals and ROIs under alfaxalone suggests that it may be better suited for fMRI studies than the co-administration of isoflurane and ketamine.

## 5. Conclusion

Anesthetics are important to many experimental approaches in animal research. However, without good, consistently applied protocols for neuroimaging, findings from these studies remain difficult to consolidate, and the degree to which they can be generalized to understand the brain in its natural neural state remains unclear. In addition to the advantages described above, there are some practical implications that favor the use of alfaxalone: 1) a single-agent protocol is easier to maintain over extended testing, is preferred by veterinary staff, and reduces concerns related to drug interactions; 2) unlike ketamine, alfaxalone in not a controlled substance, and is thus easier to obtain and store; and 3) because anesthetic depth is more consistent across the testing session under alfaxalone, the duration of testing is more predictable and concerns related to the animal waking up within the magnet are reduced. For all of these reasons, we therefore propose that alfaxalone may be a superior anesthetic agent for the safe and reliable collection of fMRI data.

## 6. Acknowledgments

We would like to thank Stephen Gordon and Alessandra Sacco for their assistance with animal care and data collection, and Trevor Szekeres for technical assistance. This research was undertaken thanks to funding from the Canada First Research Excellence Fund awarded to BrainsCAN at Western University, the Natural Sciences and Engineering Research Council of Canada, Canadian Institutes of Health Research, and the Canada Foundation for Innovation.

